# Acute inflammatory events attenuate high-sucrose diet-induced neurodegenerative processes in reproductively normal female wild-type mice

**DOI:** 10.1101/863670

**Authors:** Anthony G. Pacholko, Lane K. Bekar

## Abstract

It is known that diabetic and chronic inflammatory conditions can increase the risk of Alzheimer’s disease (AD)-like neurodegeneration in isolation. As certain elements of the diabetic/pre-diabetic state may sensitize the brain to inflammatory insult (*i.e*. excess glucocorticoid activity), there is reason to believe that obesogenic and inflammatory factors may accelerate neurodegeneration in a synergistic manner. Also, given that most AD research utilizes male animal models despite increased prevalence of AD among women, we sought to characterize elements of the established (in males) high-sucrose model of neurodegeneration, for the first time, in reproductively normal (pre-menopausal) female mice. A high-sucrose diet (20% of the drinking water) was combined with systemic intraperitoneal lipopolysaccharide (LPS) injections (0.1 mg/kg; 1x/month over 3 months) over seven months in reproductively normal female wild-type mice (C57Bl/6; n=10/group). Although a deleterious effect was hypothesized, low-dose LPS proved to protect against high sucrose diet-induced pathologies in female wild-type mice. Results from our high-sucrose group confirmed that a high-sucrose diet is a mild model of neurodegeneration in wild-type females, as evidenced by exaggerated glucocorticoid expression, spatial learning deficits, irregularities within the insulin pathway, and increased β-amyloid production and Tau phosphorylation. While LPS had little to no effect in isolation, it exerted a protective influence when added to animals sustained on a high-sucrose diet. Corticosterone homeostasis, and levels of amyloid-β (Aβ) and pTau were rescued following addition of LPS. The work presented supports a high-sucrose diet as a model of mild neurodegeneration in female mice and highlights a protective role for transient inflammation against dietary-insult that may be sex dependent.

## Introduction

Alzheimer’s disease (AD) prevalence is estimated to double by the year 2050(1,2), with approximately two-thirds of patients expected to be female, highlighting the post-menopausal loss of estrogen(3–6) as a potential risk factor. Coincident with the burgeoning AD crisis are similar trends in obesity, diabetes, and chronic inflammatory conditions, suggesting a role for lifestyle factors in disease pathogenesis. Of concern, the bulk of present dementia research relies on humanized genetic mutations that account for less than 10% of all AD cases. While these models have contributed to our understanding of disease mechanisms, they fail to elucidate the etiology of the far more prevalent sporadic late-onset variant of the disease. Furthermore, as the rate of increased incidence suggests it is unlikely for genetic factors to be sole contributors to upward trends in disease prevalence, non-transgenic animal models – particularly those dealing with environmental and/or lifestyle-mimetic stressors/risk factors – are needed to explore the pathogenesis of sporadic late-onset dementia. Also, the influence of sex on neurodegeneration must be characterized to unravel the causative factors behind increased AD incidence among women.

Excess sucrose (20% sucrose in the drinking water) has been shown (in *male* mice) to induce metabolic and pathological changes consistent with AD-related neurodegeneration in transgenic models(7–9). While only a mild phenotype is present after numerous months, multiple pathways involved in the pathogenesis of AD are altered. 1) Caloric excess is associated with elevated glucocorticoid expression(10), which has been demonstrated to quench CNS antioxidant capacity(11), potentiate neuroinflammation(12), and induce brain insulin resistance(13). Intriguingly, glucocorticoids can suppress estrogen levels and counteract estrogen activity (estrogen is neuroprotective(4,14)), suggesting a heightened sensitivity of women to dietary stressors(5). 2) Both caloric excess and chronic glucocorticoid activity can lead to liver damage and steatohepatitis(15–17), which are known to promote production of ceramides through induction of the stress-sensitive salvage pathway(18–21). Over-expression of ceramides can disrupt insulin signaling(22,23) and trigger pro-apoptotic pathways(24,25). 3) Dysregulation of brain insulin signaling, perhaps precipitated through glucocorticoid and/or ceramide overexpression, is linked to increased β-amyloidogenesis and hyperphosphorylation of the microtubule-associated Tau protein, the hallmark processes of AD pathogenesis(8,26,27). Thus, excess glucocorticoid and/or ceramide signaling/activity may link obesogenic feeding (*i.e*. high-sucrose diets) to AD-like neurodegeneration.

High sugar diets may also instigate processes that allow for the transition from acute inflammation to chronic. Caloric excess contributes to inflammation directly, thereby raising the baseline expression of inflammatory mediators and potentially allowing for inflammatory events, which may otherwise have had limited effect, to precipitate a chronic inflammatory state(28–30). Caloric excess may also enhance propagation of inflammatory mediators into the brain through disruption of the blood-brain-barrier (BBB)(31,32) and glucocorticoid-mediated alteration of anti-inflammatory and antioxidant capacities(11,12). Chronic inflammation is associated with increased β-secretase(33)- and γ-secretase activity(34), which promotes the β-amyloidogenic processing of amyloid precursor protein (APP) into Aβ. Thus, the establishment of chronic neuroinflammation is considered a potent contributor to AD-like neurodegeneration.

To explore the potential for synergistic interactions between inflammatory events and high sugar diets the present study combined, for the first time, a high-sucrose diet (20% of the drinking water) with repeated mild systemic lipopolysaccharide (LPS; 0.1 mg/kg) injections to accelerate development of neurodegeneration in a reproductively normal female wild-type mouse. We hypothesized that a high sucrose diet would enable establishment of a chronic inflammatory state in response to a low dose of LPS that is likely insufficient to drive significant neuroinflammation on its own. Reproductively normal female mice were used as a first step in the characterization of the role of estrogen in the high-sucrose model of neurodegeneration. As estrogen is known to be both anti-inflammatory and anti-β-amyloidogenic(4,14), pre-menopausal females may demonstrate a heightened resistance to neurodegeneration relative to their male and/or post-menopausal female counterparts.

## Materials and Methods

### Experimental mice and study groups

Three-month-old **female** C57/Bl6 (Charles River, Canada) mice were randomized into four groups of ten animals constituting a 2×2 design. The control group was provided normal drinking water and given three intraperitoneal (IP) saline injections delivered once per month for three months beginning after the 4^th^ week of treatment. A lipopolysaccharide (LPS; Sigma, 0.1 mg/kg IP) group was administered LPS in place of saline. A high-sucrose group was provided 20% sucrose in drinking water with three IP saline injections. A high-sucrose-LPS (hSL) combined treatment group followed the regimens of hS and LPS treated mice. See figure 1 for a summary of experimental groups and treatments (Fig. 1A). Mice were housed in pairs and kept on a 12-hr light/dark cycle. Behavioral testing began after six months of treatment with animals being sacrificed for tissues after seven months. All experiments were approved by the University of Saskatchewan Animal Research Ethics Board and done according to the Canadian Council on Animal Care. All surgery was performed under xylazine and urethane anesthesia.

**Fig 1.**
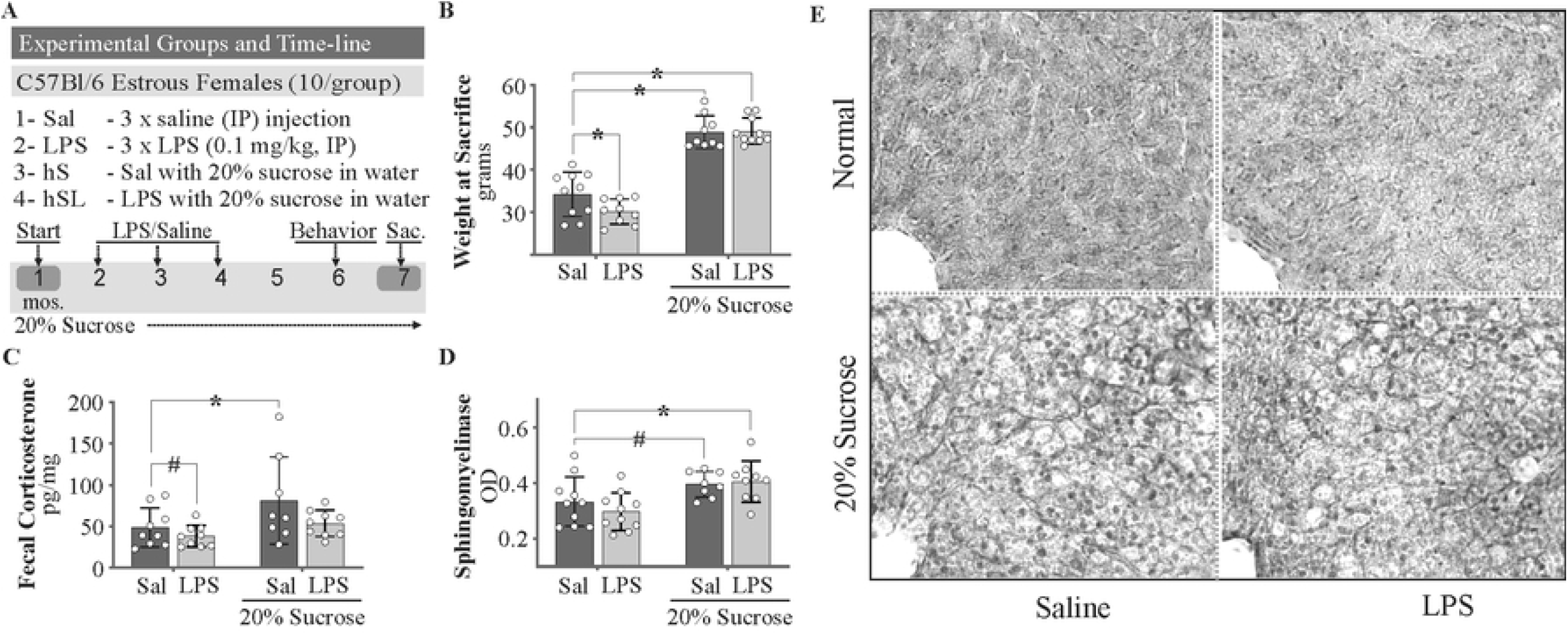
A high-sucrose model of stress and liver steatosis. **A**. Schematic outline of experimental groups and timeline. **B**. Weight gain was observed in all mice sustained on sucrose in drinking water. **C**. High sucrose diet-induced elevations in fecal corticosterone were abolished in the presence of LPS. **D**. Liver sphingomyelinase activity was increased in all sucrose treated animals. **E**. Hematoxylin and Eosin staining of 30 μm thick liver sections with the central vein apparent within a single hepatic lobule (bottom left). Lipid droplets are observed as smaller, irregular holes in the tissue. Two-way ANOVA with Fisher’s LSD post-hoc test. *P < 0.05 vs control. # Cohen’s D > 0.5. n = 8-10 for all groups.

### Behavior

After six months, mice were subjected to behavioral testing that included the open field test (OFT) for general locomotion and thigmotaxis (wall-seeking behavior) and the Barnes maze (BM) for spatial memory acquisition and retrieval. Animals were habituated to the testing room for a period of 30 minutes prior to each trial/testing. All testing was video recorded and analyzed offline using Ethovision XT 11.0 software (Leesberg, VA).

#### Open Field Test

Mice were placed in an opaque white box (35 x 35 x 30 cm) under bright white light and allowed to explore for 10 minutes (min). Animals were scored for total distance travelled and time spent within the center of the field defined as nose, body and tail >6 cm away from all walls.

#### Barnes Maze

Protocol adapted from Attar *et* al (2013)(35). The Barnes maze featured a dark escape box (placed beneath the escape hole of the stationary platform), white 100 cm (diameter) rotating platform with twenty 5.5 cm (diameter) escape holes evenly distributed about the perimeter, and a white 100 cm (diameter) stationary base with one escape location. The rotating platform was placed on top of the stationary base (itself 12 inches above the ground). ***Habituation***: mice were habituated to the maze on the first day of testing via placement in the center of the testing field (within a translucent container) for 30 seconds. Animals were then gently nudged toward the escape location over a period of 30 seconds. Upon entry, animals were habituated to the escape box for two minutes. Escape location was rotated every fourth mouse (one from each of four groups); assigned holes were held constant throughout the acquisition protocol. ***Acquisition***: mice were placed in the center of the maze under a translucent container. After ten seconds, the container was lifted, and an aversive buzzer was triggered. Animals were allowed 3 minutes to locate the escape hole under duress of the buzzer. If successful, the buzzer was silenced, and mice were held in the escape box for two minutes prior to return to their home cages. If unsuccessful, mice were guided to the escape location, followed by cessation of the buzzer and two-minute reinforcement in the escape box. This procedure was repeated once daily for seven consecutive days. Animals were scored for latency to escape (exploration of either the escape location or the holes immediately adjacent) and number of errors (exploration of incorrect holes), which were used to create an acquisition index (time to escape x # of errors averaged over the 7 days). A high acquisition index is suggestive of impaired spatial learning. ***Probe***: mice were probed for long-term recall of the escape location 48 hours after the final acquisition trial. Animals were allowed 3 minutes to explore the platform with the escape location closed. Animals were scored for latency to previous escape hole and number of errors, which were used to create a probe index (time to escape x # of errors). A high probe index suggests impaired long-term spatial recall.

### Biochemistry

After seven months, mice were weighed and sacrificed for harvesting of liver and brain tissue. One-half of the brain was flash frozen in isopentane and ground into powder for long-term storage at −80° C.

#### Acetylcholinesterase Activity Assay

Mouse brain tissue was homogenized in cold 0.1 M phosphate buffered saline (PBS; pH 8.0; 10 μl/mg) and protein content was measured by the Pierce™ Bicinchoninic Acid protein assay (BCA; Thermofisher Scientific). 5 μl (4 μg protein/μl) of the sample was added to the reaction mixture that included 300 μl of 0.1 M PBS (pH 8), 2 μl of 0.075M acetylthiocholine, and 10 μl of 0.01 M DTNB (5,5-dithiobis (2-nitro benzoic acid)) solution for a total volume of 317 μl/well. 96-well plates were measured on a spectrophotometer (SpectraMax M5, Molecular Devices) at 415 nm (25 °C) in 5 min intervals for 40 min. The maximum slopes over a 10 min period were normalized to the average control activity for final comparisons. The above protocol was derived from the original Ellman cholinesterase assay(36).

#### Nitrate/Nitrite Colorimetric Assay

Mouse brain tissue was homogenized in cold tris lysis buffer (TLB; 0.01M, pH 7.4, 3x protease/phosphatase inhibitors) and protein content was measured using the BCA assay. 55 μL of samples (4 μg protein/μL) and standards (Item No. 780014) were incubated with 10 μL each of nitrate reductase cofactors (Item No. 780012; Cayman Chemicals) and enzymes (Item No. 780010; Cayman Chemicals) for 3 hours at room temperature, followed by deproteinization with 1:1 acetonitrile. Samples/standards were vortexed for 1 minute prior to centrifugation (4° C) for 10 minutes at 10,000 rcf. Supernatants (80 μL) were placed into a 96 well plate. 50 μL of Greiss reagent 1 (Item No. 780018; Cayman Chemicals) and Greiss reagent 2 (Item No. 780020; Cayman Chemicals) were added, followed by 10-minute color development. End Point absorbance was measured on a spectrophotometer (SpectraMax M5, Molecular Devices) at 550 nm (25 °C).

#### Corticosterone ELISA

Fecal pellets were collected from the colon during sacrifice and immediately stored on dry ice for subsequent ethanol extraction (100 μL ethanol/10 mg fecal powder) and equal volume measurement of corticosterone metabolites using an ELISA-based Assay Kit (Arbor Assays). Fecal samples were chosen over serum as they better represent a long-term average less influenced by rapid stress-induced changes in corticosterone expression. It has been reported that fecal measurements reflect corticosterone levels from approximately 6-12 hours prior to defecation.

#### GSK3β/pGSK3β, Akt/pAkt, Amyloid-β40/Amyloid-β42, and Total Tau/pTau Electrochemiluminescence

Mouse brain tissue was homogenized in cold TLB (0.01M, pH 7.4, 3x protease/phosphatase inhibitors) and protein content was measured using the BCA assay (4 μg protein/μL). 20 μg of protein was assessed for total and phospho GSK3β and Akt protein concentrations, 100 μg of protein for Aβ40 and Aβ42 expression, and 10 μg of protein for total and phosphorylated Tau, as per individual Assay Kit instructions (Meso Scale Discovery). Plates were read on the MESO QuickPlex SQ 120 instrument and analyzed using the associated Workbench Discovery software (Meso Scale Discovery). Phosphorylation at Ser-9, Ser-473, and Thr-231 residues was assessed for GSK3β, Akt and Tau, respectively.

#### IRS1/pIRS1, mTOR/pmTOR and IRS2 ELISA

Mouse brain tissue was homogenized in cold TLB (0.01M, pH 7.4, 3x protease/phosphatase inhibitors) and protein content was measured using the BCA assay (diluted to 4 μg protein/μL). Tissue homogenates were further diluted to 2 μg protein/μL using TLB without protease/phosphatase inhibitors before being packaged and shipped to EVE technologies (Calgary, Canada) for ELISA-based Assay (EVE Tech). Total and phosphorylated (human Ser-636, correlates to mouse Ser-634) IRS1 and mTOR (Ser-2448) were quantified.

#### Neutral Sphingomyelinase

Mouse liver tissue was homogenized in cold TLB (0.01M, pH 7.4, 3x protease/phosphatase inhibitors) and protein content was measured using the BCA assay (diluted to 4 μg protein/μL). 200 μg of sample were used for analysis of sphingomyelinase activity via ELISA-based assay (Cayman chemicals). Kinetic and Endpoint absorbance were measured on a spectrophotometer (SpectraMax M5, Molecular Devices).

### Histology

#### Hematoxylin and Eosin Staining

Female mouse livers were fixed in neutral buffered formalin (Thermo Fisher Scientific) prior to long-term storage in PBS (0.01M, pH 7.4). Fixed liver tissues were sectioned on a vibratome (50 μm, Leica VT1200) and stained with hematoxylin and eosin for assessment of fatty liver (ballooning hepatocytes and lipid droplets). Protocol was performed according to commercially available kit instructions (Abcam assay).

### Statistical analysis

Data are expressed as mean ± SEM and compared using two-way ANOVA to assess treatment effects and potential interactions between a high-sucrose diet and repeated lipopolysaccharide challenge. Fisher’s LSD post-hoc tests were performed to assess the significance of differences between group means relative to control. Prism V 8.1.2 (GraphPad Software, Inc. SD, CA) was used to analyze two-way ANOVA data. As both the small sample size used (n=10) and the lengthy duration of the study (7 months) likely contributed to increased variation within groups, Cohen’s D measure of effect size was also used to assess the magnitude of the difference between group means(37–39). Cohen’s D values > 0.5 represent a medium effect size while values > 0.8 denote a large effect. Effect size pertains to the standard deviations by which two group means differ; *i.e*. a D of 0.5 suggests that two groups’ means differ by half a standard deviation(37). It has been proposed that small effect sizes (*x* < 0.5) are likely to be trivial, regardless of significance, whereas medium to large effects (*x* > 0.5) warrant further exploration, even in cases where statistical significance is not reached(37,40,41). A *p*-value of 0.05 is not necessarily more reliable than one of 0.08, as the difference is likely the result of minute differences in sample size (*i.e. n* of 9 vs. *n* of 10)(40). It can be argued that the size of an effect is more important than its likelihood of being attributable to type I error, so long as the associated *p*-value is reasonable. For this reason, *p*-values are provided along with the magnitude of effect (Cohen’s D). Seemingly meaningful differences between groups should not be disregarded simply-because an alpha greater than 0.05 was calculated(39,41–43). P/D_LPS_, P/D_hS_, and P/D_hSL_ pertain to the differences between the control mean and LPS, hS, and hSL group means, respectively.

## Results

### A high-sucrose diet as a model of chronic stress and liver steatosis in female wild-type mice

Perturbations in glucocorticoid homeostasis and ceramide activity have been linked to both diabetic and neurodegenerative conditions(12,13,22,23). As such, we examined measures of fecal corticosterone and liver sphingomyelinase activity to approximate the relative degree of glucocorticoid and ceramide expression. All animals sustained on 20% sucrose in their drinking water (hS) demonstrated weight gain (sucrose effect P < 0.001; D_hS_ = 3.19, D_hSL_ = 3.48; Fig. 1B) whereas LPS-treated mice showed a small decrease in weight (P/D_LPS_ = 0.030/0.96) at the time of sacrifice (7 months). Interestingly, hS-treated mice showed an increase in fecal corticosterone (sucrose effect P = 0.023) with the LPS-treated group showing a mild decrease (P/D_LPS_ = 0.46/0.55) and the hS group displaying an elevation (P/D_hS_ = 0.031/0.80; Fig. 1C) that was lost with coadministration of LPS. Increased liver neutral sphingomyelinase activity (sucrose effect P = 0.001; P/D_hS_ = 0.076/0.88, P/D_hSL_ = 0.036/0.88; Fig. 1D) and hepatic steatosis (Fig. 1E) were observed in all high sucrose fed animals (hS and hSL). Given that corticosterone injections are known to affect ceramide production and steatosis, it is interesting to find that LPS did not show any antagonistic effects on sphingomyelinase activity or steatosis in the combined group, despite normalizing fecal corticosterone levels, supporting the notion that a high sucrose diet may increase sphingomyelinase activity independent of baseline corticosterone levels.

### Seven months on a high-sucrose diet alters insulin-related signaling in female wild-type mice

As aberrant glucocorticoid and ceramide activity is associated with dysregulated brain insulin signaling(12,13,22,23), we examined baseline total and phospho concentrations of insulin pathway-associated second messenger protein analytes in hemibrain homogenates to probe for any irregularities. It should be noted that the second messengers explored are not exclusive to the insulin pathway. It is therefore possible that any differences observed might not be solely attributable to insulin dysfunction.

Total insulin-receptor substrate 1 (IRS1) protein levels were increased after seven months in all mice on a 20% sucrose diet (sucrose effect P < 0.001; P/D_hS_ = 0.007/1.57, P/D_hSL_ = 0.056/0.924; Fig. 2A), while phosphorylated levels remained relatively consistent; hS-treated mice showing a mild increase (P/D_hS_ = 0.342/0.710; Fig. 2B). Suggesting reduced feedback inhibition, the ratio of phosphorylated IRS1 to total IRS1 showed a mild decrease in high sucrose fed mice (sucrose effect P = 0.140; P/D_hS_ = 0.190/0.747, P/D_hSL_ = 0.276/0.553; Fig. 2C), perhaps on account of decreased pathway activity. Conversely, activation of the downstream mediator Akt was increased following LPS treatment, regardless of the presence or absence of a high sucrose diet (LPS effect P = 0.010; P/D_LPS_ = 0.028/1.11, P/D_hSL_ = 0.226/0.73; Fig. 2D). No increase in the phosphorylation state of Akt (Fig 2D), mTOR (Fig. 2E), or GSK3β (Fig. 2F) – downstream mediators of the insulin pathway – were noted in high-sucrose fed mice. In fact, levels of *p*mTOR were reduced in all high sucrose fed mice, suggesting suppressed activity (sucrose effect P = 0.007; P/D_hS_ = 0.118/0.819, P/D_hSL_ = 0.032/1.20; Fig. 2E). This lack of second messenger activation downstream of IRS1 despite increased availability of the protein may suggest an impairment or dysregulation induced by hS, but not LPS, within the brain insulin pathway.

**Fig 2.**
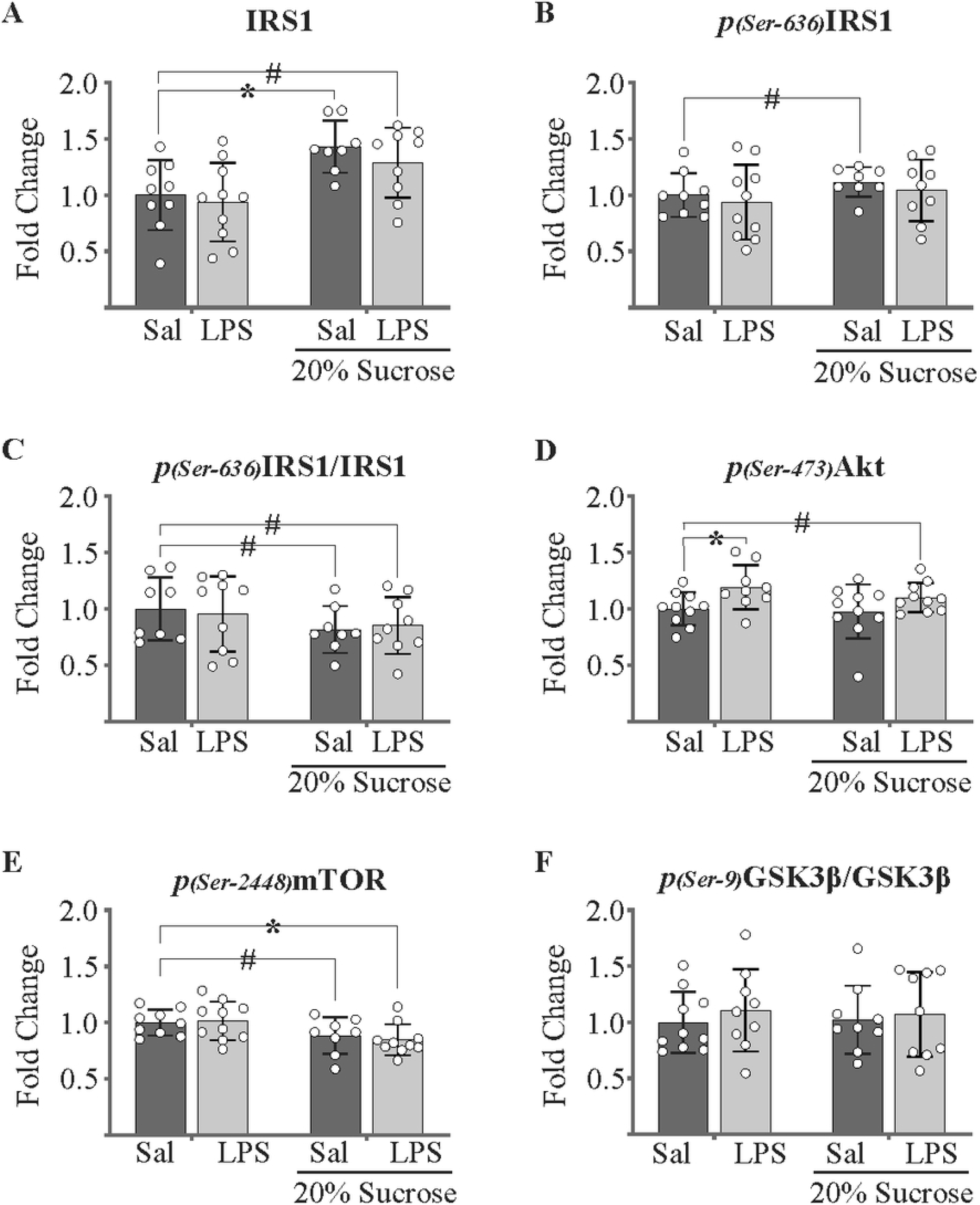
Seven months on a high-sucrose diet promotes early irregularities in brain insulin pathway activity. **A.** High sucrose treatment increased total IRS1 protein levels. **B**. Phospho-IRS1 levels were only mildly affected. **C**. A reduced degree of IRS1 phosphorylation was observed. **D**. Akt phosphorylation was elevated in LPS treated animals, regardless of the presence/absence of sucrose. **E**. Phosphorylation of mTOR was suppressed in all mice receiving sucrose in the drinking water. **F**. No changes in GSK3β phosphorylation were observed. All values are normalized to control (fold change for y-axis). Two-way ANOVA with Fisher’s LSD post-hoc test. *P < 0.05 vs control. # Cohen’s D > 0.5. n = 9-10 for all groups.

### High-sucrose treatment contributes to neurodegenerative processes in female wild-type mice

Considering the well-established role of neuroinflammation in the etiology and progression of neurodegenerative conditions, we examined hemibrain homogenate nitric oxide activity and cytokine expression to assess any potential neuroinflammatory phenotype. Furthermore, given the importance of acetylcholine in memory and cognition, and its depleted state in numerous neurodegenerative conditions, we evaluated acetylcholinesterase activity. Nitrate + nitrite levels, indicative of NO activity, were found to be elevated in all groups receiving high sucrose in the drinking water (sucrose effect P = 0.002; P/D_hS_ = 0.027/0.932; P/D_hSL_ < 0.001/1.742) as well as mildly elevated in the LPS-treated group (P/D_LPS_ = 0.227/0.537, Fig. 3A). No changes in acetylcholinesterase activity were observed in any group (Fig. 3B). Although the anti-inflammatory IL-10 and inflammatory TNF-α, IL-1β, IL-5 and IL-6 showed a few differences between groups (Table 1), when data were normalized against IL-10 (no changes observed across all groups) variation was improved (Fig. 3C-F). Interestingly, although all inflammatory cytokines showed a sucrose effect except for IL-6, only the hS diet with concurrent LPS treatment resulted in a significant reduction in levels of the proinflammatory cytokines TNF-α (sucrose effect P = 0.040; P/D_hSL_ = 0.093/1.21), IL-1β (sucrose effect P = 0.021; P/D_hSL_ = 0.021/1.23), IL-5 (sucrose effect P = 0.014; P/D_hSL_ = 0.012/1.27) and IL-6 (sucrose effect P = 0.288; P/D_hSL_ = 0.078/0.928).

**Fig 3.**
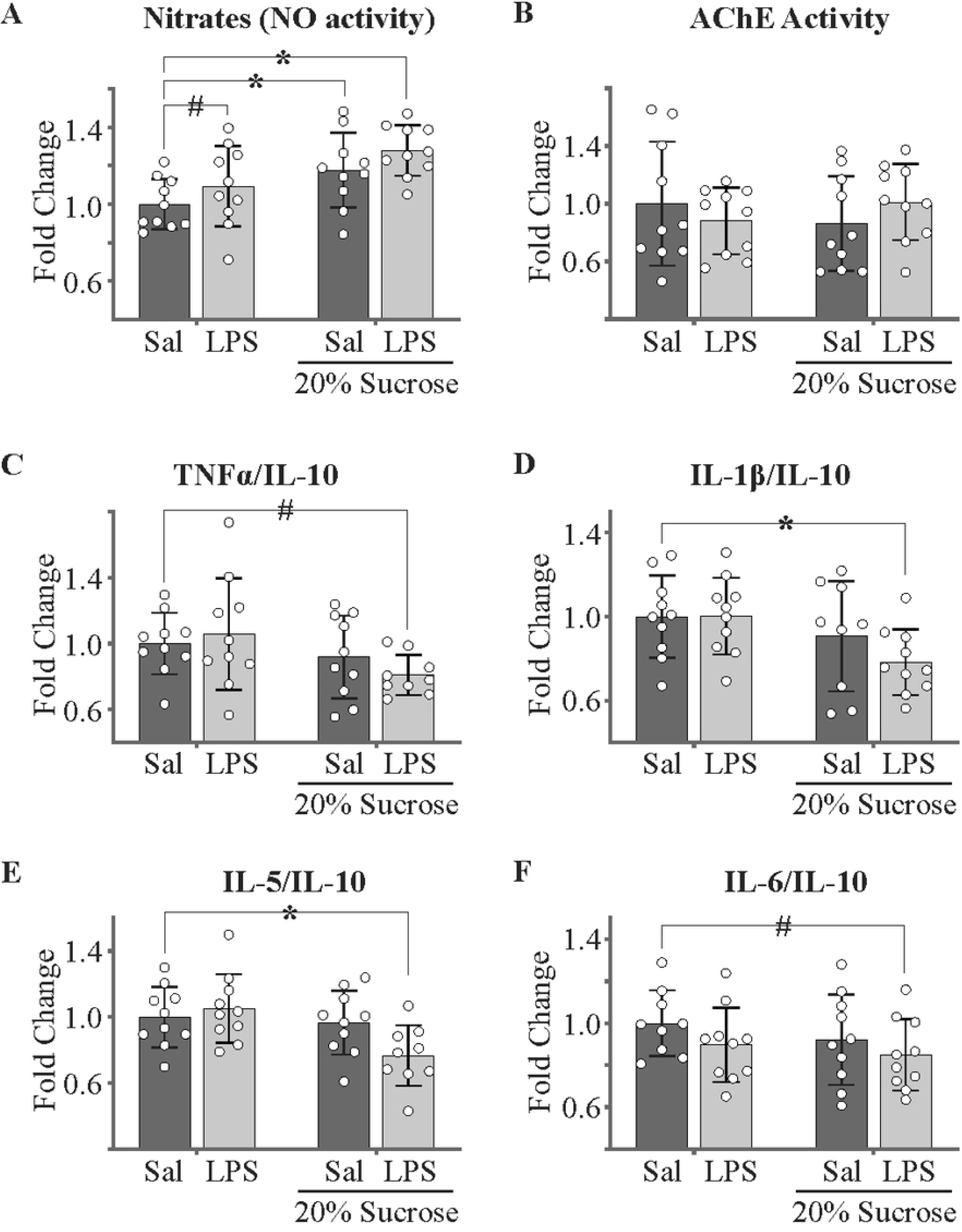
A high-sucrose diet and repeated lipopolysaccharide challenge exert differing effects on general inflammation. **A**. A high sucrose diet increased nitrate + nitrite expression independent of LPS exposure. **B**. No notable effects on acetylcholinesterase activity were observed. **C-F**. Only the combined high sucrose and LPS treated group showed a decrease in inflammatory cytokines. Cytokines were normalized to the anti-inflammatory cytokine IL-10 to help reduce variation within groups. All values are normalized to control (fold change for y-axis). Two-way ANOVA with Fisher’s LSD post-hoc test. *P < 0.05 vs control. # Cohen’s D > 0.5. n = 9-10 for all groups.

**Table 1.** Hemibrain cytokine expression (pg/ml). Open box means Cohen’s D > 0.5. Grey box indicates P < 0.05 via two-way ANOVA Fisher’s LSD. N = 9-10 for all groups.

|  | <u>IL-10</u> | <u>TNF-α</u> | <u>IL-1β</u> | <u>IL-5</u> | <u>IL-6</u> |
| --- | --- | --- | --- | --- | --- |
| **Con** | 2.71 ± 0.39 | 0.32 ± 0.07 | 0.85 ± 0.22 | 0.33 ± 0.06 | 6.99 ± 0.69 |
| **LPS** | 2.79 ± 0.56 | 0.34 ± 0.11 | 0.86 ± 0.16 | 0.35 ± 0.06 | 6.38 ± 0.83 |
| **hS** | 2.81 ± 0.66 | 0.29 ± 0.07 | 0.76 ± 0.20 | 0.33 ± 0.06 | 6.56 ± 1.09 |
| **hSL** | 2.90 ± 0.51 | 0.28 ± 0.07 | 0.70 ± 0.12 | 0.26 ± 0.06 | 6.31 ± 0.54 |

As both hyper- and hypo-activation of the insulin pathway are known to contribute to increased genesis of the amyloid-β_42_ protein (hallmarks of dementia-associated neurodegeneration)(18,44,45) and hyperphosphorylation of the microtubule associated protein Tau, we assessed treatment effects on hemibrain Aβ_42_ and Tau levels. Mice in all treatment groups displayed an increasing effect on Aβ_42_ production compared to control (P/D_LPS_ = 0.077/0.776; P/D_hS_ = 0.059/0.783; P/D_hSL_ = 0.185/0.601, Fig. 4A, top). Only a mild effect on Aβ_40_ levels was observed in the combined group (P/D_hSL_ = 0.177/0.560, Fig. 4A, middle). While both LPS (P/D_LPS_ = 0.030/0.946) and hS (P/D_hS_ = 0.085/0.701; Fig. 4A, bottom) treatment increased the ratio of Aβ_42_ to Aβ_40_ proteins individually, this effect was lost in combination, supporting an antagonistic interaction (interaction P = 0.013, Fig. 4A, bottom). Only a mild decrease of total Tau expression was observed in the combined group (P/D_hSL_ = 0.357/0.520). Treatment with LPS suppressed Tau phosphorylation, regardless of the presence or absence of high sucrose (LPS effect P = 0.006; P/D_LPS_ = 0.022/1.14; P/D_hSL_ = 0.015/1.18, Fig. 4B middle), though this effect was lost when normalized to total Tau protein (Fig. 3B bottom).

**Fig 4.**
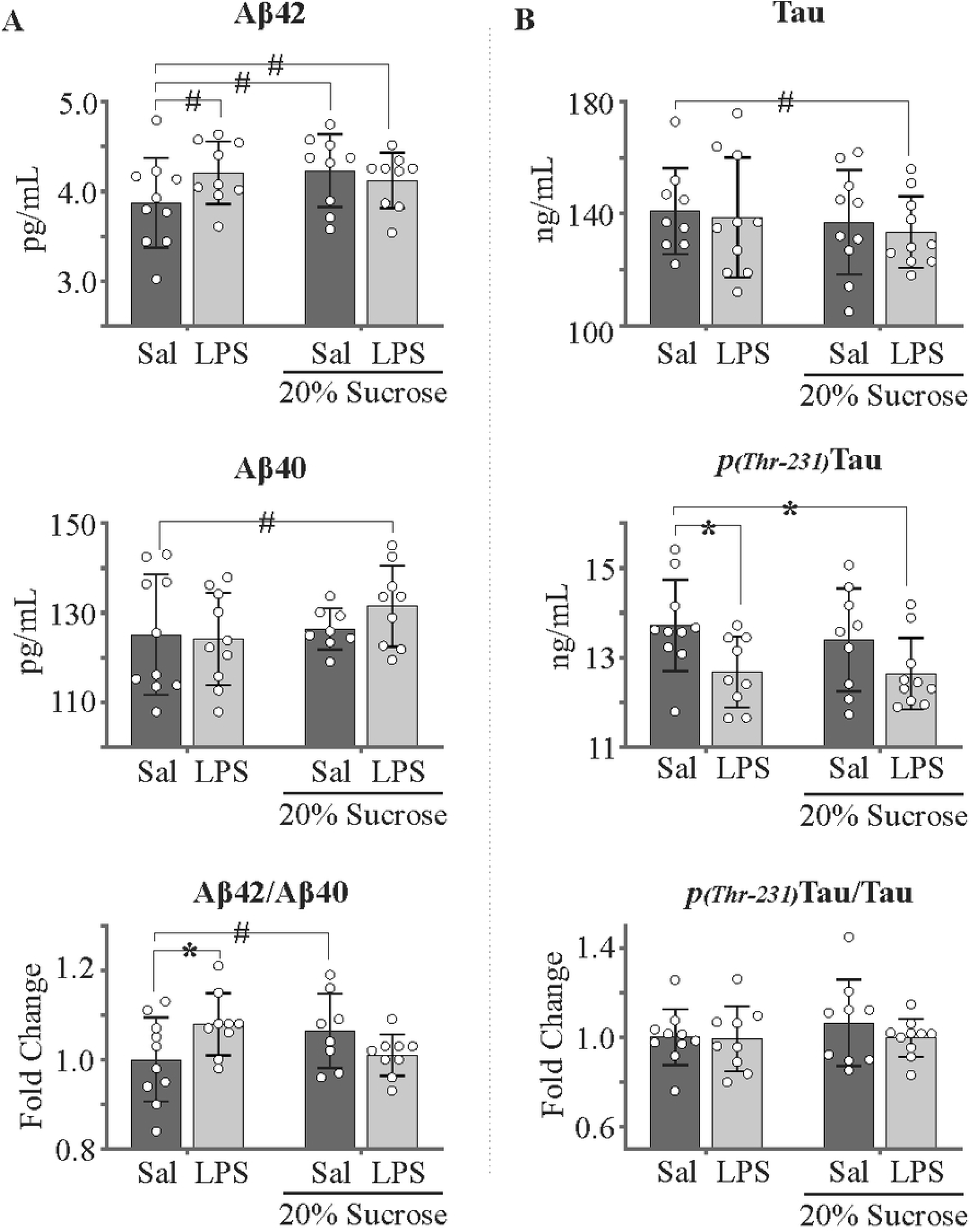
Effects of a high-sucrose diet and/or lipopolysaccharide on β-amyloidogenesis and Tau phosphorylation in reproductively normal female mice. **A.** Aβ_42_ expression demonstrated an elevation in response to both LPS treatment and high-sucrose diet. Aβ_40_ was only affected in the combined treatment group. **B**. Although only the combined treatment affected total tau (top), LPS had a significant effect on phosphotau (middle). High sucrose in the water had a small effect (Cohen’s D = 0.39) on the ratio of phosphotau over total tau. Two-way ANOVA with Fisher’s LSD post-hoc test. *P < 0.05 vs control. # Cohen’s D > 0.5. n = 8-10 for all groups.

### A high sugar diet alters behavior differently in the presence/absence of lipopolysaccharide

AD-like neurodegeneration often manifests phenotypically as memory impairments and behavioral abnormalities. Thus, we employed simple Barnes maze and Open Field protocols to assess spatial memory and anxiety-like behavior, respectively. Behavioral trials performed during the 6^th^ month of treatment demonstrated differing high-sucrose-associated behavioral phenotypes in the presence and absence of lipopolysaccharide. Animals sustained on hS alone displayed the largest impairment to spatial learning performance in the Barnes maze, characterized by increased latency to escape and frequency of errors (P/D_hS_ = 0.040/1.01; Fig. 5A, left). Such results are consistent with the effects of chronic corticosterone on spatial learning observed elsewhere(9,46,47), suggesting a glucocorticoid-mediated impairment in our hS mice. In contradiction to this notion, hSL mice demonstrated a mild impairment in spatial learning (P = 0.127; D_hSL_ = 0.842, Fig. 5A, left) that did not coincide with increased fecal corticosterone levels. Interestingly, although LPS or hS treatment had no effect on long-term spatial recall alone, when combined we observed an improvement in these reproductively normal female mice (Fig. 5A, right).

**Fig 5.**
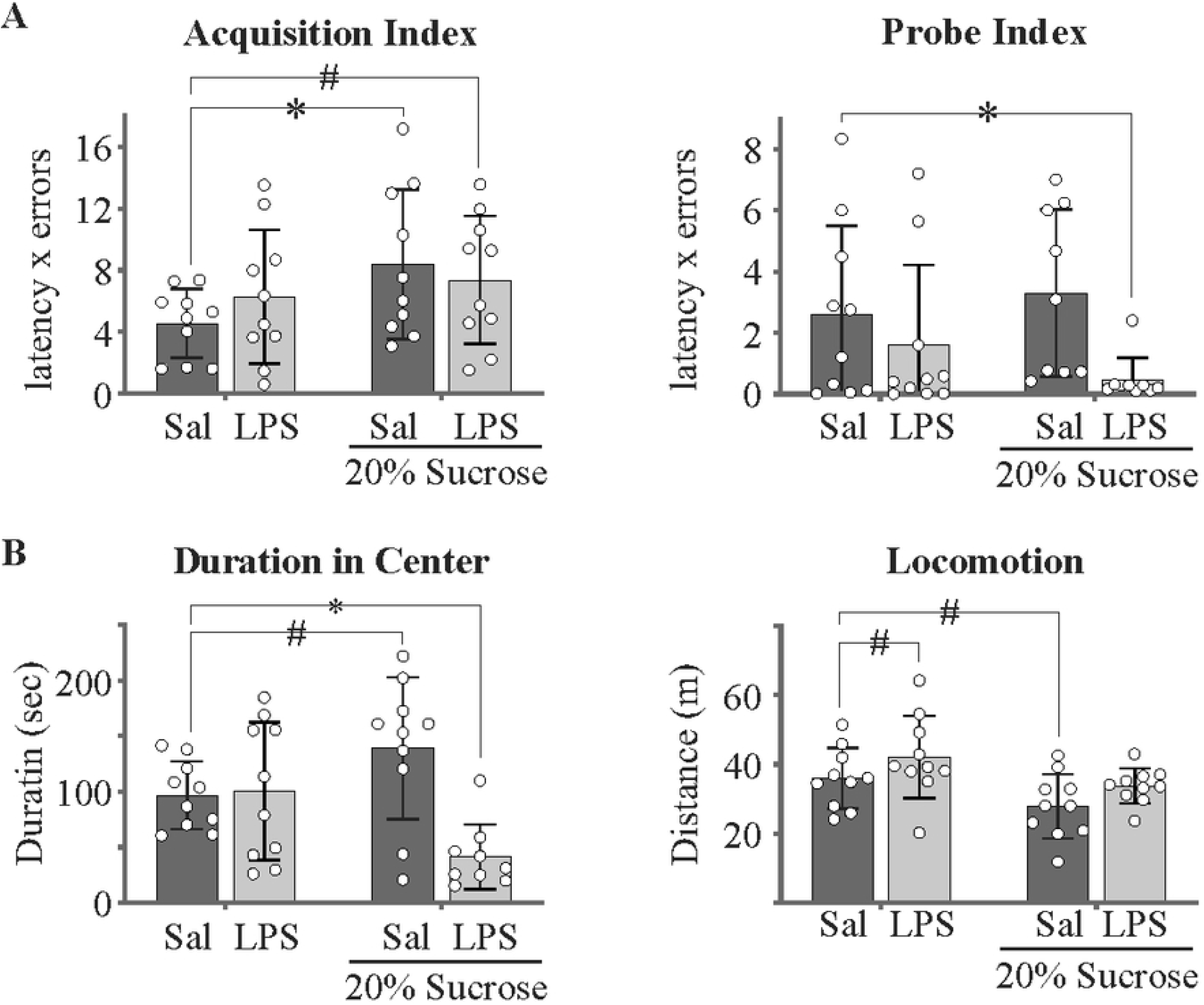
Differential alteration of Barnes maze-associated spatial learning performance and Open Field degree of thigmotaxis in response to high-sucrose feeding and/or systemic LPS injection. **A**. Rate of spatial memory acquisition was impaired in all animals sustained on a high-sucrose diet, though the aaddition of LPS attenuated - but did not abolish - these deficits. Only the combined treatment group showed an improvement in 48 hour recall of the escape location. **B**. The addition of systemic LPS to high-sucrose fed mice appeared to promote anxiety-like behavior (increased thigmotaxis) in the open field. Animals fed a high sugar diet demonstrated a trend toward increased duration in center. Locomotion did not differ significantly between groups, though a trend toward increased and reduced distance traveled was noted for LPS and hS animals, respectively. Two-way ANOVA with Fisher’s LSD post-hoc test. *P < 0.05 vs control. # Cohen’s D > 0.5. n = 10 for all groups.

During the Open Field test, there was an LPS-mediated effect that was only observed as a decrease of time spent in the center in the combined group (LPS effect P = 0.006, P/D_hSL_ = 0.021/1.85; Fig. 5B, left). Although no sucrose effect was observed, the hS group showed a mild increase in time spent in the center (P/D_hS_ = 0.063/0.854). Avoidance of the center of the field is often associated with anxiety-like behavior, suggesting that an anxiogenic interaction (interaction P = 0.003, Fig. 5B, left) between hS and LPS exists that is absent following the individual treatments. While hS-treatment reduced locomotion (sucrose effect P = 0.007) in the Open Field, both LPS (P/D_LPS_ = 0.138/0.591) and hS (P/D_hS_ = 0.056/0.891) had mild effects on locomotion that were lost in the combined group (Fig. 5B right). The lack of effect in the combined group suggests effects on locomotion do not account for the increased anxiety-like behavior observed in this group.

## Discussion

Given that trends in AD incidence coincide with similar trends in obesity, diabetes, and chronic inflammation - all known risk factors for AD – the present study combined a high-sucrose drinking water regimen with repeated monthly intraperitoneal lipopolysaccharide injections to accelerate AD-related pathology in estrous-female wild-type mice. Female mice were used as the bulk of AD literature concerns male animal models despite nearly two-thirds of all AD patients being women(2). We demonstrate that high sugar consumption promotes upregulation of early-stage AD-related processes after just 6-7 months, as evidenced by elevated fecal corticosterone (Fig. 1C), increased liver sphingomyelinase activity (Fig. 1D), brain insulin pathway dysregulation (Fig. 2), increased β-amyloidogenesis (Fig. 4A), an altered brain inflammation state (Fig. 3 and Table 1), and worsened spatial learning in the Barnes maze (Fig. 5). Interestingly, the addition of LPS prevented/lessened many of these effects; fecal corticosterone (Fig. 1C) was normal, Tau phosphorylation (Fig. 4B) was suppressed, and spatial acquisition deficits were curtailed.

Previous studies have shown that caloric excess promotes lipogenesis and triglyceride storage in both the liver and adipose tissue(48). In cases of chronic excess, exacerbated intake leads to liver insulin resistance and steatohepatitis (inflammation of fatty liver)(15,16,49). This inflammatory state promotes lipolysis and degeneration of the liver, culminating in mitochondrial- and/or apoptotic-mediated cell-death and ceramide synthesis(50,51). The resultant free fatty acids, proinflammatory cytokines, and ceramides can induce both systemic and central dysregulation of the insulin pathway, leading to eventual neurotoxicity(16,18). In this study, animals sustained on 20% sucrose in their drinking water displayed weight gain, hepatic steatosis, elevated fecal corticosterone expression, and elevated liver neutral sphingomyelinase activity (ceramides) after 7 months. As noted prior, these conditions have been linked with insulin resistance, neurodegeneration and cognitive impairment(16,19,52,53). A recent study by Chen, *et al* (2017) demonstrated that liver ceramide synthesis is upregulated by glucocorticoid signaling(53), thus opening the possibility that glucocorticoids are partly responsible for the high-sucrose-induced liver pathologies in this study. While we did not assess systemic insulin sensitivity directly, observed increases in sphingomyelinase activity and liver steatosis suggest that our high-sucrose animals were likely insulin resistant. Ceramides have been shown to inhibit both peripheral and central insulin signaling(18,52,53), and to promote neuroinflammation(54) and oxidative stress in the CNS(16,19,52). Total IRS1 proteins were found to be increased in whole hemi-brain homogenates of high sucrose fed mice without concomitant phosphorylation of downstream targets such as Akt and mTOR, suggesting a potential compensatory increase in IRS1 proteins in response to reduced insulin signaling. Although glucocorticoids are known to induce brain insulin resistance(13), observed irregularities within the brain insulin pathway appear to be attributable to ceramide-mediated mechanisms. Irregularities were displayed even in high sucrose fed mice with normal glucocorticoid activity (hSL group), implicating ceramides as the likely driving force behind the observed dysregulation. Furthermore, elevations in total IRS1 proteins without concurrent increase in Akt phosphorylation are consistent with results observed following injection with ceramides, as demonstrated by de la Monte, *et al* (2010)(52). It should be noted that the second messengers discussed are not exclusive to the insulin pathway. Both insulin-like growth factor 1 (IGF1)(55)- and estrogen-receptor (ER)(56–58)-mediated signaling incorporate elements of the PI3K-Akt-mTOR pathway, and thus present potential confounds. As the mice used in this study are female, the interpretations regarding ceramides and insulin pathway dysfunction should be viewed within the context of these limitations: any differences in mTOR or Akt phosphorylation could have been influenced by estrogen, and therefore may not represent insulin resistance.

We previously demonstrated that a 20% sucrose diet increases Tau phosphorylation after just 4-months in *male* mice(9). Similar – though lesser – observations were noted in the present work, confirming for the first time the high-sucrose diet as a model of mild neurodegeneration in *female* wild-type mice. Both total Aβ_42_ and the ratio of Aβ_42_ to Aβ_40_ were increased by the high sugar diet. Furthermore, high-sucrose animals displayed increased nitrate + nitrite expression, suggesting enhanced NO production (perhaps through upregulated iNOS). Exaggerated NO has been linked to neuroinflammation and nitrosative activity, both of which are known to enhance the processing of APP to Aβ(33,34). Increased β-amyloidogenesis and NO expression were associated with worsened spatial learning performance, highlighting a potential decline in cognition. Given the data presented, it seems possible that a high-sugar diet upregulated neurodegenerative processes (*i.e*. β-amyloidogenesis, nitrosative activity, etc.) through glucocorticoid(13)- and hepatic ceramide-mediated mechanisms(16,19,19) to influence spatial memory.

It should be noted that the mild phenotype observed may be related to the sex of the animals. Estrogen has been shown to exert neuroprotection in a variety of models (cell culture as well as animal) and can diminish many pathological processes associated with AD, such as β-amyloidopathy, glucocorticoid over-expression, mitochondrial dysfunction, and oxidative stress(4,5). Also, the mild elevation in Aβ_42_ coupled with impaired spatial memory acquisition but intact long-term recall could represent an early time-point in late-onset AD pathogenesis(59–62) (Fig. 6), at least in females. Given the proposed neuroprotective effects of estrogen against the AD-related pathological processes associated with insulin dysregulation – such as inhibition of GSK3(56), suppression of tau phosphorylation(56,63), and attenuation of Aβ production(64) – it is possible that male and/or post-menopausal female mice would display more severe neurodegenerative phenotypes under the experimental conditions employed herein; *i.e*. employing similar treatment parameters, we previously demonstrated a more robust phenotype after just 4 months in male mice. Regardless, the results presented in this study using reproductively normal females assist in the establishment of an important baseline for future exploration of the role of estrogen in dietary-stressor-induced neurodegeneration.

**Fig 6.**
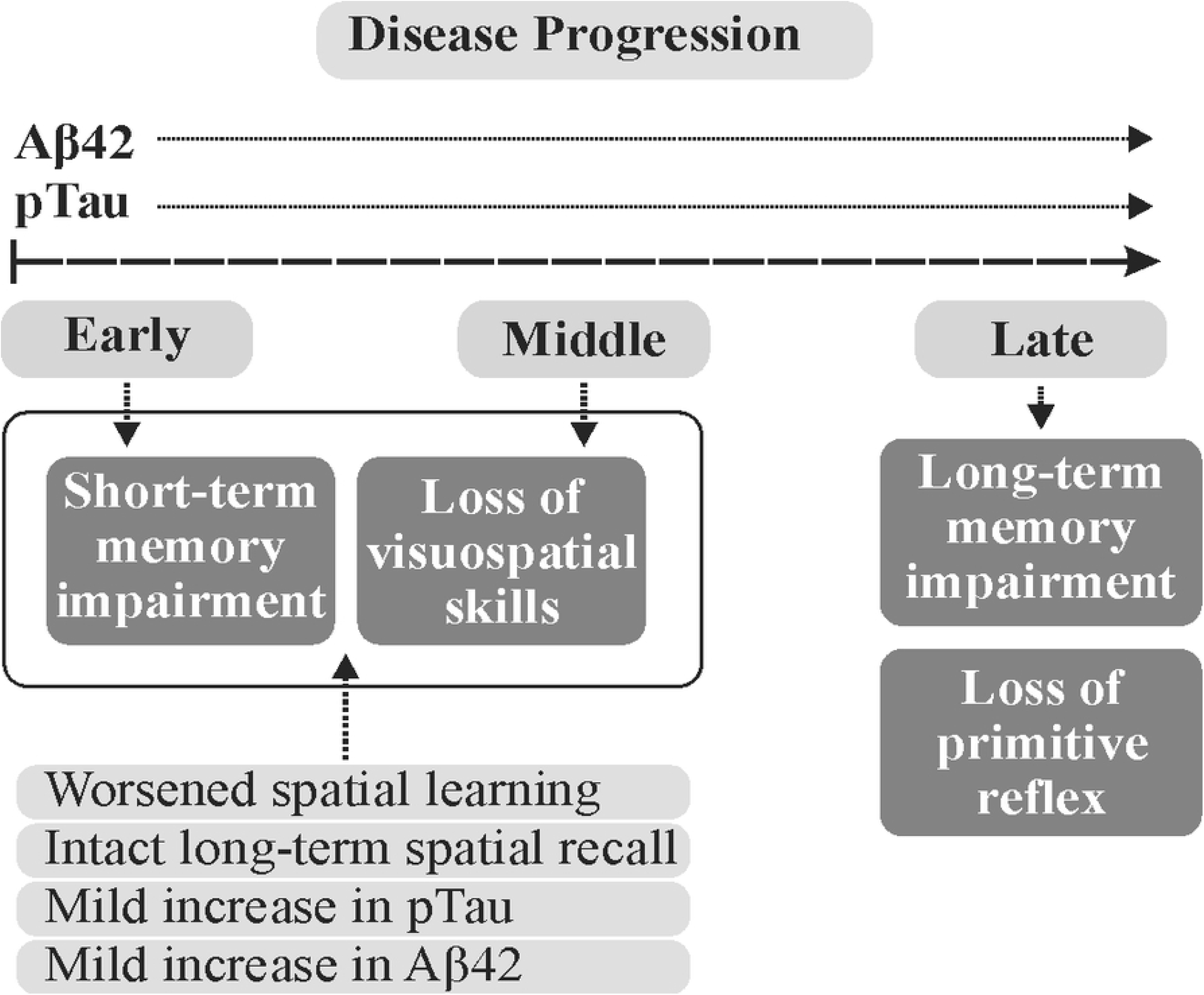
Seven months on a high-sucrose diet induces behavioral abnormalities and upregulation of neurodegenerative processes suggestive of an early late-onset AD-like phenotype. Early AD is characterized, in part, by impairments in short-term memory formation. As the disease progresses, long-term recall begins to diminish. This coincides with increased Aβ and pTau burden. Our mice demonstrate a mild though present Aβ and pTau phenotype that coincides with worsened spatial acquisition (delayed learning of a visual escape location in the Barnes maze) and spared long-term recall. This may suggest that our mice display an AD-like phenotype representative of an early time-point within late-onset pathogenesis.

The modest effects of hS treatment on Tau phosphorylation and β-amyloidogenesis could also have been due to the use of hemibrain homogenates, which may have diluted region-specific differences. In addition, it has been proposed that while diabetes alone may be insufficient to generate a robust AD phenotype, it could serve as a cofactor in the etiology and progression of the disease(49). Nonetheless, when paired with corticosterone, sphingomyelinase, Aβ, and pTau data, the results obtained in a non-transgenic wild-type model at this early time-point support a high-sucrose diet-mediated upregulation of neurodegenerative processes.

Counter to our expectations that LPS would exacerbate the effects of a high-sugar diet, its addition failed to accelerate development of a neurodegenerative phenotype. Instead, the combination appeared to be protective, as antagonistic interactions were noted at several levels. First and perhaps most notably, animals on the combined high-sucrose and LPS regimen did not display an elevation in fecal corticosterone. Given the proposed central nature of glucocorticoids to high-sucrose diet-mediated pathology(13), attenuation of corticosterone expression may have proved to be quite beneficial. In fact, acute LPS-mediated inflammation was found to suppress Tau phosphorylation (with or without sucrose) in animals sustained on a high sugar diet. Interestingly, the only pathology that arose due to the combination was observed in the Open Field, where the addition of LPS to high-sucrose animals enhanced thigmotaxis. Avoidance of the center has been linked to anxiety-like behavior, suggesting the combination treatment to be anxiogenic.

The protective effects of LPS may have been due to the transient nature of the inflammation induced. Glucocorticoids engage in a form of negative feedback through a hippocampal-HPA axis circuit. Once a temporal and spatial threshold of interaction between glucocorticoids and their receptors in the hippocampus is reached, activation of the HPA is suppressed. This process allows for robust yet transient corticosterone/cortisol responses to stressful stimuli(65). Acute inflammation has been shown to promote a robust increase in circulating glucocorticoids(66) capable of triggering negative feedback. Thus, it is possible that this spike in corticosterone activity may have ‘reset’ the high-sucrose diet-induced chronic elevation in glucocorticoids through activation of the aforementioned feedback loop (Fig. 7).

**Fig 7.**
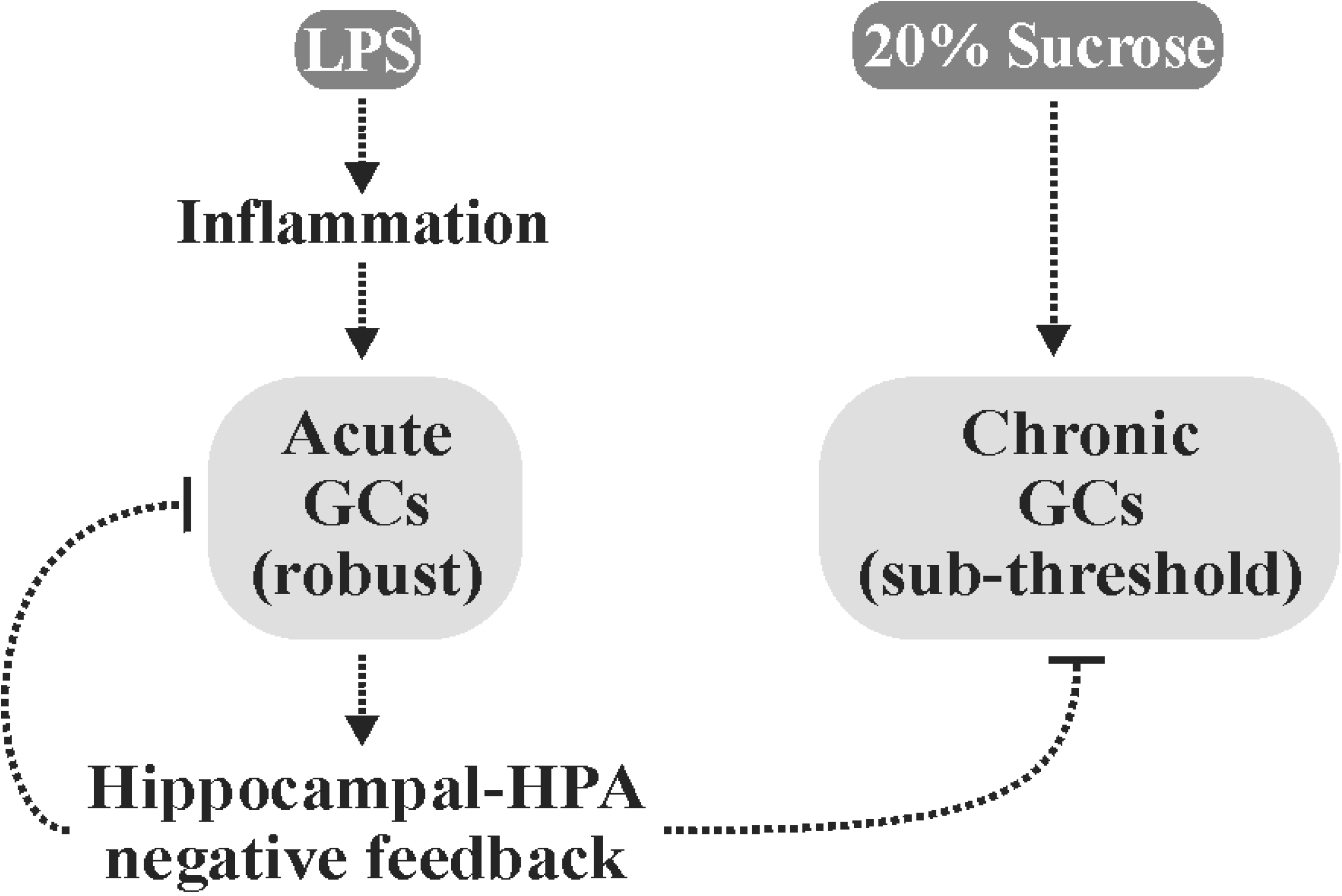
LPS may have conferred protection via attenuation of chronic glucocorticoid signaling. LPS is known to induce a robust inflammatory response associated with elevated glucocorticoid release via activation of the HPA axis. Glucocorticoids self-regulate their activity through induction of negative feedback via a hippocampal-HPA axis-dependent loop. High-sucrose diets could potentially promote chronic increases in glucocorticoid expression without triggering this negative feedback (sub-threshold elevations). Transient and robust glucocorticoid responses following LPS administration may have ‘re-set’ this chronic increase.

LPS demonstrates dose-specific effects, with high doses leading to sepsis(67); however, the effects at low concentrations are far more controversial. While it has been reported that sustained treatment with low-dose LPS promotes low-grade inflammation(68), transient exposure has been found to result in suppressed inflammatory responses to future challenge(69–71). Short-term exposure to low-dose LPS has even been shown to suppress inflammation and prevent neurodegeneration following cerebral ischemia-reperfusion injury(72–74). With regard to the present study, it is important to note that the final injection of LPS occurred four months prior to sacrifice, making it difficult to compare the neuroinflammation results herein with the literature. The addition of low-dose LPS to our hS-fed mice suppressed proinflammatory cytokine expression (IL-5, IL-6 and IL-1β), suggesting a possible state of inflammatory hypo-responsiveness caused by low-dose LPS-induced tolerance to TLR4 ligands(69–71). Alternatively, Everhardt and associates (2016) observed that female mice injected with LPS display subdued proinflammatory responses relative to males(14), which raises the possibility that the lack of LPS-induced neuroinflammation observed in this study could be attributable to estrogen interference. Furthermore, gonadectomy leads to an increase in mortality in female rats exposed to LPS that is attenuated by estrogen replacement therapy(75). Taking potential estrogen interference into account, it is possible that male and/or post-menopausal females would demonstrate more robust neuroinflammatory phenotypes in response to the dose of LPS administered in this study.

If transient low-dose LPS-induced inflammation did protect against select elements of the deleterious effects of a high-sucrose diet by ‘re-setting’ the corticosterone response in these reproductively normal female mice, it would reinforce the notion that high sugar diet-induced pathology is mediated primarily by glucocorticoids, and that transient inflammatory events may be beneficial in the long-term management of chronic stress.

## Conclusion

This study supports the efficacy of high sugar diets in the induction of pathologic processes involved in neurodegeneration and demonstrates that transient inflammatory events may antagonize pathologies associated with said diets in reproductively normal female mice. While it remains possible that modern lifestyle changes and inflammatory medical complications can intersect to accelerate development of an AD phenotype, it remains clear that much work is to be done to untangle the relationships between diet, stress and neurodegeneration. Furthermore, this study may highlight a potential reason as to why AD occurrence varies among individuals. If acute stress/inflammation is protective in females, then perhaps differences in lifestyles, diet, exercise and work/home environments contribute to long-term cognitive health; *i.e*. stress is needed to ‘re-set’ cortisol homeostasis. It also remains likely that males, females and post-menopausal females respond differently to inflammatory events on account of differing sex hormone profiles, thus raising the possibility that the combination of a high sugar diet and LPS injection would be neurotoxic in males and/or post-menopausal females.

## Acknowledgments

Funding: This work was supported by the Saskatchewan Health Research Foundation (grant # 3075) and University of Saskatchewan (College of Medicine Research Award).

## Conflicts of Interest

The authors declare that there are no conflicts of interest.

## References

1. Dementia in Saskatchewan | Alzheimer Society of Saskatchewan [Internet]. [cited 2019 Mar 30]. Available from: https://alzheimer.ca/en/sk/Vote-Dementia-Care/Dementia-SK

2. Alzheimer’s Association. 2016 Alzheimer’s disease facts and figures. Alzheimers Dement J Alzheimers Assoc. 2016 Apr;12(4):459–509.

3. Manly JJ, Merchant CA, Jacobs DM, Small SA, Bell K, Ferin M, et al. Endogenous estrogen levels and Alzheimer’s disease among postmenopausal women. Neurology. 2000 Feb 22;54(4):833–7.

4. Pike CJ, Carroll JC, Rosario ER, Barron AM. Protective actions of sex steroid hormones in Alzheimer’s disease. Front Neuroendocrinol. 2009 Jul;30(2):239–58.

5. Herrera AY, Mather M. Actions and interactions of estradiol and glucocorticoids in cognition and the brain: Implications for aging women. Neurosci Biobehav Rev. 2015 Aug;55:36–52.

6. Rosario ER, Chang L, Head EH, Stanczyk FZ, Pike CJ. Brain levels of sex steroid hormones in men and women during normal aging and in Alzheimer’s disease. Neurobiol Aging. 2011 Apr;32(4):604–13.

7. Carvalho C, Machado N, Mota PC, Correia SC, Cardoso S, Santos RX, et al. Type 2 diabetic and Alzheimer’s disease mice present similar behavioral, cognitive, and vascular anomalies. J Alzheimers Dis JAD. 2013;35(3):623–35.

8. Carvalho C, Cardoso S, Correia SC, Santos RX, Santos MS, Baldeiras I, et al. Metabolic alterations induced by sucrose intake and Alzheimer’s disease promote similar brain mitochondrial abnormalities. Diabetes. 2012 May;61(5):1234–42.

9. Choudhary P, Pacholko AG, Palaschuk J, Bekar LK. The locus coeruleus neurotoxin, DSP4, and/or a high sugar diet induce behavioral and biochemical alterations in wild-type mice consistent with Alzheimers related pathology. Metab Brain Dis. 2018;33(5):1563–71.

10. Stranahan AM, Arumugam TV, Cutler RG, Lee K, Egan JM, Mattson MP. Diabetes impairs hippocampal function through glucocorticoid-mediated effects on new and mature neurons. Nat Neurosci. 2008 Mar;11(3):309–17.

11. McIntosh LJ, Hong KE, Sapolsky RM. Glucocorticoids may alter antioxidant enzyme capacity in the brain: baseline studies. Brain Res. 1998 Apr 27;791(1-2):209–14.

12. Frank MG, Miguel ZD, Watkins LR, Maier SF. Prior exposure to glucocorticoids sensitizes the neuroinflammatory and peripheral inflammatory responses to E. coli lipopolysaccharide. Brain Behav Immun. 2010 Jan;24(1):19–30.

13. Osmanovic J, Plaschke K, Salkovic-Petrisic M, Grünblatt E, Riederer P, Hoyer S. Chronic exogenous corticosterone administration generates an insulin-resistant brain state in rats. Stress Amst Neth. 2010 Mar;13(2):123–31.

14. Everhardt Queen A, Moerdyk-Schauwecker M, McKee LM, Leamy LJ, Huet YM. Differential Expression of Inflammatory Cytokines and Stress Genes in Male and Female Mice in Response to a Lipopolysaccharide Challenge. PloS One. 2016;11(4):e0152289.

15. Capeau J. Insulin resistance and steatosis in humans. Diabetes Metab. 2008 Dec;34(6 Pt 2):649–57.

16. Lyn-Cook LE, Lawton M, Tong M, Silbermann E, Longato L, Jiao P, et al. Hepatic Ceramide May Mediate Brain Insulin Resistance and Neurodegeneration in Type 2 Diabetes and Non-alcoholic Steatohepatitis. J Alzheimers Dis JAD. 2009 Apr;16(4):715–29.

17. Lieber CS, Leo MA, Mak KM, Xu Y, Cao Q, Ren C, et al. Model of nonalcoholic steatohepatitis. Am J Clin Nutr. 2004 Mar;79(3):502–9.

18. Monte DL, M S. Triangulated Mal-Signaling in Alzheimer’s Disease: Roles of Neurotoxic Ceramides, ER Stress, and Insulin Resistance Reviewed. J Alzheimers Dis. 2012 Jan 1;30(s2):S231–49.

19. de la Monte SM. Metabolic derangements mediate cognitive impairment and Alzheimer’s disease: role of peripheral insulin resistance diseases. Panminerva Med. 2012 Sep;54(3):171–8.

20. Summers SA. Ceramides in insulin resistance and lipotoxicity. Prog Lipid Res. 2006 Jan;45(1):42–72.

21. Turpin SM, Nicholls HT, Willmes DM, Mourier A, Brodesser S, Wunderlich CM, et al. Obesity-Induced CerS6-Dependent C16:0 Ceramide Production Promotes Weight Gain and Glucose Intolerance. Cell Metab. 2014 Oct 7;20(4):678–86.

22. Bourbon NA, Sandirasegarane L, Kester M. Ceramide-induced inhibition of Akt is mediated through protein kinase Czeta: implications for growth arrest. J Biol Chem. 2002 Feb 1;277(5):3286–92.

23. Chalfant CE, Kishikawa K, Mumby MC, Kamibayashi C, Bielawska A, Hannun YA. Long chain ceramides activate protein phosphatase-1 and protein phosphatase-2A. Activation is stereospecific and regulated by phosphatidic acid. J Biol Chem. 1999 Jul 16;274(29):20313–7.

24. Woodcock J. Sphingosine and ceramide signalling in apoptosis. IUBMB Life. 2006 Aug;58(8):462–6.

25. Cuvillier O, Andrieu-Abadie N, Ségui B, Malagarie-Cazenave S, Tardy C, Bonhoure E, et al. [Sphingolipid-mediated apoptotic signaling pathways]. J Soc Biol. 2003;197(3):217–21.

26. Cao D, Lu H, Lewis TL, Li L. Intake of sucrose-sweetened water induces insulin resistance and exacerbates memory deficits and amyloidosis in a transgenic mouse model of Alzheimer disease. J Biol Chem. 2007 Dec 14;282(50):36275–82.

27. Correia SC, Santos RX, Perry G, Zhu X, Moreira PI, Smith MA. Insulin-resistant brain state: the culprit in sporadic Alzheimer’s disease? Ageing Res Rev. 2011 Apr;10(2):264–73.

28. Shimobayashi M, Albert V, Woelnerhanssen B, Frei IC, Weissenberger D, Meyer-Gerspach AC, et al. Insulin resistance causes inflammation in adipose tissue. J Clin Invest. 2018 Apr 2;128(4):1538–50.

29. Wellen KE, Hotamisligil GS. Inflammation, stress, and diabetes. J Clin Invest. 2005 May;115(5):1111–9.

30. Burchfield JG, Kebede MA, Meoli CC, Stöckli J, Whitworth PT, Wright AL, et al. High dietary fat and sucrose result in an extensive and time-dependent deterioration in health of multiple physiological systems in mice. J Biol Chem. 2018 Apr 13;293(15):5731–45.

31. Davidson TL, Monnot A, Neal AU, Martin AA, Horton JJ, Zheng W. The effects of a high-energy diet on hippocampal-dependent discrimination performance and blood-brain barrier integrity differ for diet-induced obese and diet-resistant rats. Physiol Behav. 2012 Aug 20;107(1):26–33.

32. Kanoski SE, Zhang Y, Zheng W, Davidson TL. The effects of a high-energy diet on hippocampal function and blood-brain barrier integrity in the rat. J Alzheimers Dis JAD. 2010;21(1):207–19.

33. Kwak Y-D, Wang R, Li JJ, Zhang Y-W, Xu H, Liao F-F. Differential regulation of BACE1 expression by oxidative and nitrosative signals. Mol Neurodegener. 2011 Mar 3;6:17.

34. Guix FX, Wahle T, Vennekens K, Snellinx A, Chávez-Gutiérrez L, Ill-Raga G, et al. Modification of γ-secretase by nitrosative stress links neuronal ageing to sporadic Alzheimer’s disease. EMBO Mol Med. 2012 Jul;4(7):660–73.

35. Attar A, Liu T, Chan W-TC, Hayes J, Nejad M, Lei K, et al. A Shortened Barnes Maze Protocol Reveals Memory Deficits at 4-Months of Age in the Triple-Transgenic Mouse Model of Alzheimer’s Disease. PLoS ONE [Internet]. 2013 Nov 13 [cited 2019 Sep 10];8(11). Available from: https://www.ncbi.nlm.nih.gov/pmc/articles/PMC3827415/

36. Ellman GL, Courtney KD, Andres V, Feather-Stone RM. A new and rapid colorimetric determination of acetylcholinesterase activity. Biochem Pharmacol. 1961 Jul;7:88–95.

37. Sullivan GM, Feinn R. Using Effect Size—or Why the P Value Is Not Enough. J Grad Med Educ. 2012 Sep;4(3):279–82.

38. Bartolucci AA, Tendera M, Howard G. Meta-analysis of multiple primary prevention trials of cardiovascular events using aspirin. Am J Cardiol. 2011 Jun 15;107(12):1796–801.

39. Nahm FS. What the P values really tell us. Korean J Pain. 2017 Oct;30(4):241–2.

40. Halsey Lewis G. The reign of the p-value is over: what alternative analyses could we employ to fill the power vacuum? Biol Lett. 2019 May 31;15(5):20190174.

41. Wasserstein RL, Schirm AL, Lazar NA. Moving to a World Beyond “p < 0.05.” Am Stat. 2019 Mar 29;73(sup1):1–19.

42. Hurlbert SH, Levine RA, Utts J. Coup de Grâce for a Tough Old Bull: “Statistically Significant” Expires. Am Stat. 2019 Mar 29;73(sup1):352–7.

43. Amrhein V, Greenland S, McShane B. Scientists rise up against statistical significance. Nature. 2019 Mar;567(7748):305–7.

44. Stöhr O, Schilbach K, Moll L, Hettich MM, Freude S, Wunderlich FT, et al. Insulin receptor signaling mediates APP processing and β-amyloid accumulation without altering survival in a transgenic mouse model of Alzheimer’s disease. Age Dordr Neth. 2013 Feb;35(1):83–101.

45. Steen E, Terry BM, Rivera EJ, Cannon JL, Neely TR, Tavares R, et al. Impaired insulin and insulin-like growth factor expression and signaling mechanisms in Alzheimer’s disease--is this type 3 diabetes? J Alzheimers Dis JAD. 2005 Feb;7(1):63–80.

46. Wright RL, Lightner EN, Harman JS, Meijer OC, Conrad C D. Attenuating corticosterone levels on the day of memory assessment prevents chronic stress-induced impairments in spatial memory. Eur J Neurosci. 2006 Jul;24(2):595–605.

47. Yau JL, Olsson T, Morris RG, Meaney MJ, Seckl JR. Glucocorticoids, hippocampal corticosteroid receptor gene expression and antidepressant treatment: relationship with spatial learning in young and aged rats. Neuroscience. 1995 Jun;66(3):571–81.

48. Leonard BL, Watson RN, Loomes KM, Phillips ARJ, Cooper GJ. Insulin resistance in the Zucker diabetic fatty rat: a metabolic characterisation of obese and lean phenotypes. Acta Diabetol. 2005 Dec;42(4):162–70.

49. de la Monte SM, Wands JR. Alzheimer’s Disease Is Type 3 Diabetes–Evidence Reviewed. J Diabetes Sci Technol Online. 2008 Nov;2(6):1101–13.

50. Langeveld M, Aerts JMFG. Glycosphingolipids and insulin resistance. Prog Lipid Res. 2009 Jul;48(3-4):196–205.

51. Holland WL, Summers SA. Sphingolipids, insulin resistance, and metabolic disease: new insights from in vivo manipulation of sphingolipid metabolism. Endocr Rev. 2008 Jun;29(4):381–402.

52. de la Monte SM, Tong M, Nguyen V, Setshedi M, Longato L, Wands JR. Ceramide-mediated insulin resistance and impairment of cognitive-motor functions. J Alzheimers Dis JAD. 2010;21(3):967–84.

53. Chen T-C, Benjamin DI, Kuo T, Lee RA, Li M-L, Mar DJ, et al. The glucocorticoid-Angptl4-ceramide axis induces insulin resistance through PP2A and PKCζ. Sci Signal. 2017 Jul 25;10(489):eaai7905.

54. Scheiblich H, Schlütter A, Golenbock DT, Latz E, Martinez-Martinez P, Heneka MT. Activation of the NLRP3 inflammasome in microglia: the role of ceramide. J Neurochem. 2017;143(5):534–50.

55. Wrigley S, Arafa D, Tropea D. Insulin-Like Growth Factor 1: At the Crossroads of Brain Development and Aging. Front Cell Neurosci [Internet]. 2017 Feb 1 [cited 2019 Sep 10];11. Available from: https://www.ncbi.nlm.nih.gov/pmc/articles/PMC5285390/

56. Cardona-Gomez P, Perez M, Avila J, Garcia-Segura LM, Wandosell F. Estradiol inhibits GSK3 and regulates interaction of estrogen receptors, GSK3, and beta-catenin in the hippocampus. Mol Cell Neurosci. 2004 Mar;25(3):363–73.

57. Mendez P, Wandosell F, Garcia-Segura LM. Cross-talk between estrogen receptors and insulin-like growth factor-I receptor in the brain: cellular and molecular mechanisms. Front Neuroendocrinol. 2006 Dec;27(4):391–403.

58. Varea O, Escoll M, Diez H, Garrido JJ, Wandosell F. Oestradiol signalling through the Akt-mTORC1-S6K1. Biochim Biophys Acta. 2013 May;1833(5):1052–64.

59. Kumar A, Tsao JW. Alzheimer Disease. In: StatPearls [Internet]. Treasure Island (FL): StatPearls Publishing; 2019 [cited 2019 Apr 8]. Available from: http://www.ncbi.nlm.nih.gov/books/NBK499922/

60. Liljegren M, Landqvist Waldö M, Rydbeck R, Englund E. Police Interactions Among Neuropathologically Confirmed Dementia Patients: Prevalence and Cause. Alzheimer Dis Assoc Disord. 2018 Dec;32(4):346–50.

61. Petersen RC. How early can we diagnose Alzheimer disease (and is it sufficient)? The 2017 Wartenberg lecture. Neurology. 2018 Aug 28;91(9):395–402.

62. Schachter AS, Davis KL. Alzheimer’s disease. Dialogues Clin Neurosci. 2000 Jun;2(2):91–100.

63. Carroll JC, Rosario ER. The potential use of hormone-based therapeutics for the treatment of Alzheimer’s disease. Curr Alzheimer Res. 2012 Jan;9(1):18–34.

64. Levin-Allerhand JA, Lominska CE, Wang J, Smith JD. 17Alpha-estradiol and 17beta-estradiol treatments are effective in lowering cerebral amyloid-beta levels in AbetaPPSWE transgenic mice. J Alzheimers Dis JAD. 2002 Dec;4(6):449–57.

65. Sterner EY, Kalynchuk LE. Behavioral and neurobiological consequences of prolonged glucocorticoid exposure in rats: relevance to depression. Prog Neuropsychopharmacol Biol Psychiatry. 2010 Jun 30;34(5):777–90.

66. Basharat S, Parker JA, Murphy KG, Bloom SR, Buckingham JC, John CD. Leptin fails to blunt the lipopolysaccharide-induced activation of the hypothalamic-pituitary-adrenal axis in rats. J Endocrinol. 2014 May;221(2):229–34.

67. Opal SM. Endotoxins and other sepsis triggers. Contrib Nephrol. 2010;167:14–24.

68. Guo H, Diao N, Yuan R, Chen K, Geng S, Li M, et al. Subclinical-Dose Endotoxin Sustains Low-Grade Inflammation and Exacerbates Steatohepatitis in High-Fat Diet–Fed Mice. J Immunol. 2016 Mar 1;196(5):2300–8.

69. Chae BS. Pretreatment of Low-Dose and Super-Low-Dose LPS on the Production of In Vitro LPS-Induced Inflammatory Mediators. Toxicol Res. 2018 Jan;34(1):65–73.

70. de Vos AF, Pater JM, van den Pangaart PS, de Kruif MD, van ‘t Veer C, van der Poll T. In vivo lipopolysaccharide exposure of human blood leukocytes induces cross-tolerance to multiple TLR ligands. J Immunol Baltim Md 1950. 2009 Jul 1;183(1):533–42.

71. Pardon MC. Lipopolysaccharide hyporesponsiveness: protective or damaging response to the brain? Romanian J Morphol Embryol Rev Roum Morphol Embryol. 2015;56(3):903–13.

72. Delayre-Orthez C, de Blay F, Frossard N, Pons F. Dose-dependent effects of endotoxins on allergen sensitization and challenge in the mouse. Clin Exp Allergy J Br Soc Allergy Clin Immunol. 2004 Nov;34(11):1789–95.

73. Gong G, Huang Y, Yuan L, Hu L, Cai L. [Requirement of LRG in endotoxin-mediated brain protection]. Xi Bao Yu Fen Zi Mian Yi Xue Za Zhi Chin J Cell Mol Immunol. 2011 Aug;27(8):865–7.

74. He F, Zhang N, Lv Y, Sun W, Chen H. Low-dose lipopolysaccharide inhibits neuronal apoptosis induced by cerebral ischemia/reperfusion injury via the PI3K/Akt/FoxO1 signaling pathway in rats. Mol Med Rep. 2019 Mar 1;19(3):1443–52.

75. Merkel SM, Alexander S, Zufall E, Oliver JD, Huet-Hudson YM. Essential role for estrogen in protection against Vibrio vulnificus-induced endotoxic shock. Infect Immun. 2001 Oct;69(10):6119–22.

